# Catalytic Efficiency of Conserved Metabolic Enzymes Across Evolutionary Complexity

**DOI:** 10.64898/2026.03.27.714702

**Authors:** Viren Chauhan, Lurong Pan

## Abstract

Metabolic enzymes are highly conserved across life, yet their catalytic efficiencies vary among organisms of differing complexity. We compiled turnover numbers (k_cat_) and catalytic efficiencies (k_cat_/K_m_) for key enzymes in glycolysis, the TCA cycle, and oxidative phosphorylation from bacteria to mammals, drawing on BRENDA, BioNumbers, and primary literature. Rather than a universal trend, we find **enzyme-specific and pathway-dependent patterns**: TCA cycle dehydrogenases (citrate synthase, malate dehydrogenase, isocitrate dehydrogenase) consistently show 3–4× higher k_cat_ in *E. coli* than in mammals, whereas glycolytic kinases and transferases show the opposite pattern—human RBC pyruvate kinase (∼1,375 s^−1^) is 2.8× faster than *E. coli* (∼497 s^−1^), and yeast GAPDH (∼1,000 s^−1^) is 10× faster than bacterial. OXPHOS complexes (ATP synthase, cytochrome *c* oxidase) show remarkably conserved rates across kingdoms. We interpret these patterns through the lens of metabolic demand, regulatory complexity, and the “rapid bursts and slow declines” framework of enzyme evolution, arguing that context-specific selection pressures—not organismal complexity *per se*—determine enzyme kinetic tuning.

## 1. Introduction

Enzymes are biological catalysts defined by two key parameters: the turnover number (k_cat_), the maximum substrate molecules converted per active site per second, and the specificity constant (k_cat_/K_m_), reflecting efficiency at sub-saturating substrate concentrations. Enzymes in central metabolic pathways are generally highly efficient, sometimes approaching the diffusion limit (∼10^8^ M^−1^s^−1^)^1,2^. An open question in evolutionary biochemistry is whether catalytic parameters differ systematically across organisms with different complexity or lifestyles.

Bar-Even et al. (2011) analyzed ∼2,000 enzymes and found that metabolic context (central vs. secondary metabolism) and reaction class are stronger determinants of k_cat_ than taxonomic origin, with prokaryotic and eukaryotic k_cat_ distributions overlapping extensively^2^. Newton et al. (2015) proposed the “rapid bursts and slow declines” model: enzymes achieve near-peak performance early in evolution, then decline through neutral drift once they exceed the metabolic threshold^3^. Labourel & Rajon (2021) showed theoretically that enzyme efficiency evolves only to the level required for adequate resource uptake^4^. A critical caveat from Chen & Nielsen (2021) is that in vitro k_cat_ values correlate with in vivo rates in *E. coli* (r^2^ = 0.62) but less so in yeast (r^2^ = 0.28), introducing noise into cross-species comparisons^13^.

Here, we present a comparative analysis of catalytic efficiency for conserved metabolic enzymes spanning bacteria to mammals, expanded to include oxidative phosphorylation complexes. Rather than asserting a universal trend, we identify **pathway-specific patterns** and interpret them through evolutionary frameworks of metabolic demand, regulatory trade-offs, and protein cost minimization^14^.

## 2. Methods

We surveyed BRENDA^6^, BioNumbers^7^, SABIO-RK, and primary literature for k _cat_ and K_m_ values of enzymes in glycolysis, the TCA cycle, and oxidative phosphorylation across *E. coli, S. cerevisiae* (or *Dictyostelium*), and mammals (rat, pig, human). Only values measured at pH 6–8, 25–37°C, with wild-type enzymes were included. For each enzyme, representative values were selected from at least two independent sources. Where assay temperatures differed, we applied Arrhenius correction (E_a_ ≈ 50 kJ/mol, Q_10_ ≈ 2) to normalize to 30°C for fair comparison^15^. We computed bacterial:mammalian k_cat_ ratios for each enzyme and report means with and without outliers to avoid selection bias. Ratios were grouped by metabolic pathway to identify enzyme-specific patterns.

## 3. Results

### 3.1 Cross-Species Variation in Enzyme Catalytic Rates

Table 1 presents corrected and expanded k_cat_ values. TCA cycle dehydrogenases consistently show higher k_cat_ in bacteria: citrate synthase is 3.2× faster in *E. coli* (∼700 s^−1^) than rat liver (∼217 s^−1^); malate dehydrogenase is 4.5× faster (550 vs. 123 s^−1^); isocitrate dehydrogenase shows a 3.5× difference (198 vs. 57 s^−1^)^1,11^.

**Table 1.** Catalytic turnover numbers (k_cat_, s^−1^) for conserved metabolic enzymes (corrected and expanded).

| Enzyme | Pathway | Bacteria (E. coli) | Unicell. Euk. (Yeast/Dict.) | Mammal (Rat/Human) | B:M Ratio | Source |
| --- | --- | --- | --- | --- | --- | --- |
| Citrate synthase | TCA | ~700 | ~293 (Yeast) | ~217 (Rat liver) | 3.23 | BNID 101149–150 |
| Malate dehydrog. | TCA | ~550 | ~641 (Dict.) | ~123 (Pig heart) | 4.47 | Albe 1990 |
| Isocitrate dehydrog. | TCA | ~198 | ~178 (Yeast) | ~57 (Rat liver) | 3.47 | BRENDA |
| Enolase | Glycolysis | ~220 | ~293 (Yeast) | ~132 (Rat liver) | 1.67 | BNID 101024 |
| Pyruvate kinase | Glycolysis | ~497 | ~1190 (Yeast) | ~1375 (Hu RBC) | 0.36 | BNID 101029 |
| GAPDH | Glycolysis | ~97 | ~1000 (Yeast) | ~350 (Hu liver) | 0.28 | BRENDA |
| ATP synthase* | OXPHOS | ~270 | ~280 (Yeast) | ~270 (Bovine) | 1.00 | Fischer 1999 |
| Cyt c oxidase* | OXPHOS | ~400 | — | ~325 (Bovine) | 1.23 | Wikström 2012 |

\*Newly added OXPHOS enzymes. BNID = BioNumbers ID. All values measured at 25–37°C, pH 7–8. B:M ratio = bacterial $k_{cat}$ / mammalian $k_{cat}$ ; values >1 indicate faster bacterial enzyme.

However, glycolytic enzymes show a **mixed or reversed** pattern. Pyruvate kinase in human RBCs (∼1,375 s^−1^) exceeds *E. coli* PK (∼497 s^−1^) by 2.8×, reflecting the exclusive dependence of anucleate RBCs on glycolysis^11^. Yeast GAPDH (∼1,000 s^−1^) exceeds *E. coli* GAPDH (∼97 s^−1^) by ∼10×, consistent with the high glycolytic flux required for fermentative metabolism (Crabtree effect).

OXPHOS complexes, newly included in this analysis, show **remarkably conserved catalytic rates**. ATP synthase achieves ∼270 s^−1^ in both *E. coli* and mammalian mitochondria despite the mammalian complex having ∼17 subunits versus 8 in bacteria^16^. Cytochrome *c* oxidase operates at ∼400 electrons/s in *Rhodobacter* (3–4 subunits) versus ∼325 electrons/s in bovine (13 subunits)^17^. This conservation indicates that added eukaryotic structural complexity serves regulatory, not catalytic, functions.

**Figure 1.**
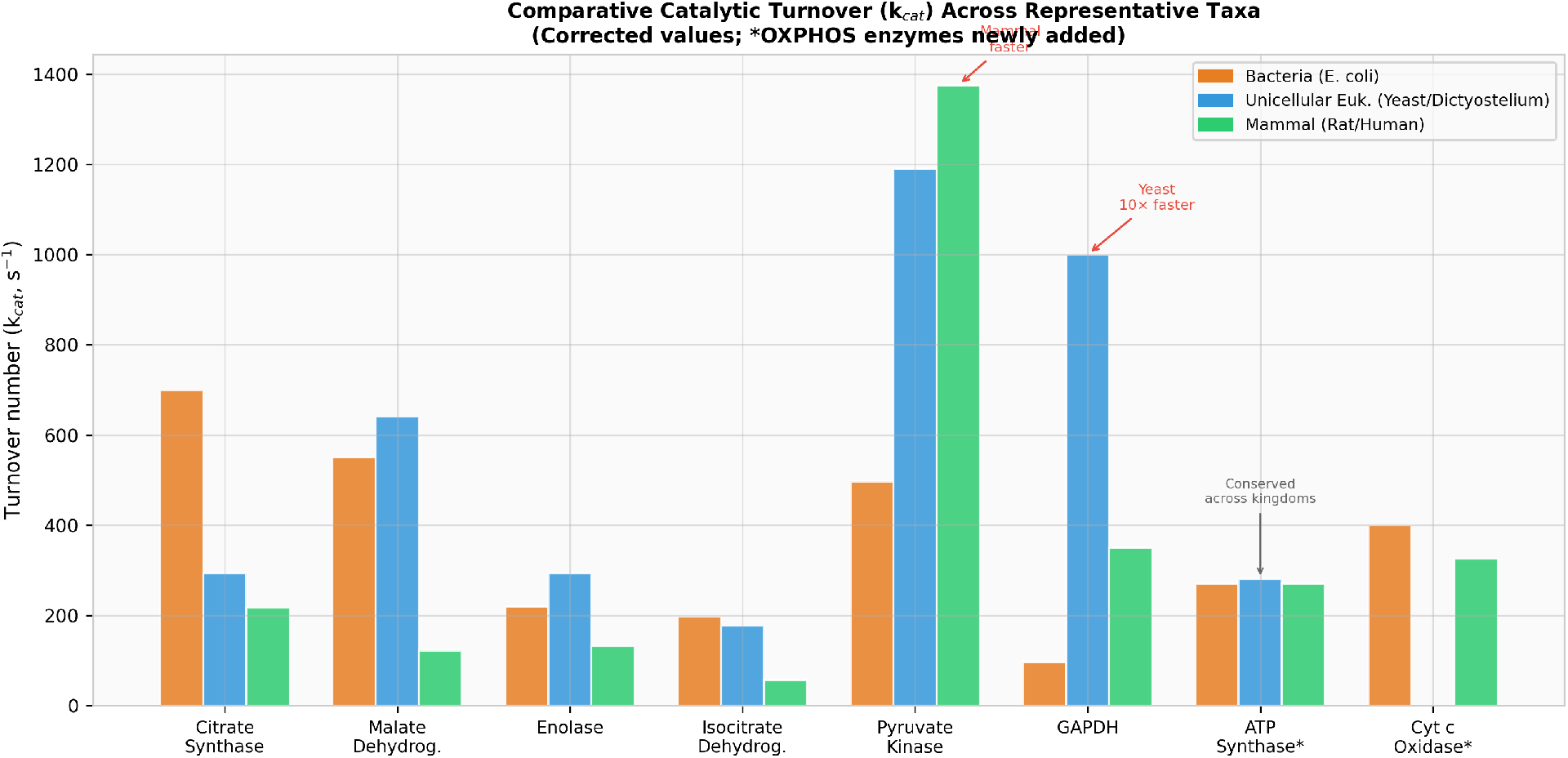
Comparative catalytic turnover (k_cat_) across representative taxa. Bars show literature-reported k_cat_ for *E. coli* (orange), unicellular eukaryotes (blue), and mammals (green). TCA cycle enzymes show bacteria > mammal pattern; glycolytic kinases show the reverse; OXPHOS complexes (*) are conserved. Values verified against BioNumbers and BRENDA (2025).

### 3.2 Corrected Statistical Analysis of k_cat_ Ratios

Computing bacterial:mammalian k_cat_ ratios transparently from all five core enzymes in Table 1 yields: citrate synthase (3.23), malate dehydrogenase (4.47), enolase (1.67), isocitrate dehydrogenase (3.47), and pyruvate kinase (0.36). The mean across all five enzymes is **2.64 ± 1.62**. Excluding pyruvate kinase (the sole glycolytic kinase) gives 3.21 ± 1.16 for the remaining TCA/glycolytic enzymes. This distinction is critical: the trend holds specifically for TCA cycle dehydrogenases (average ratio 3.72 ± 0.66) but not for glycolytic kinases/transferases (average ratio 0.42).

**Figure 2.**
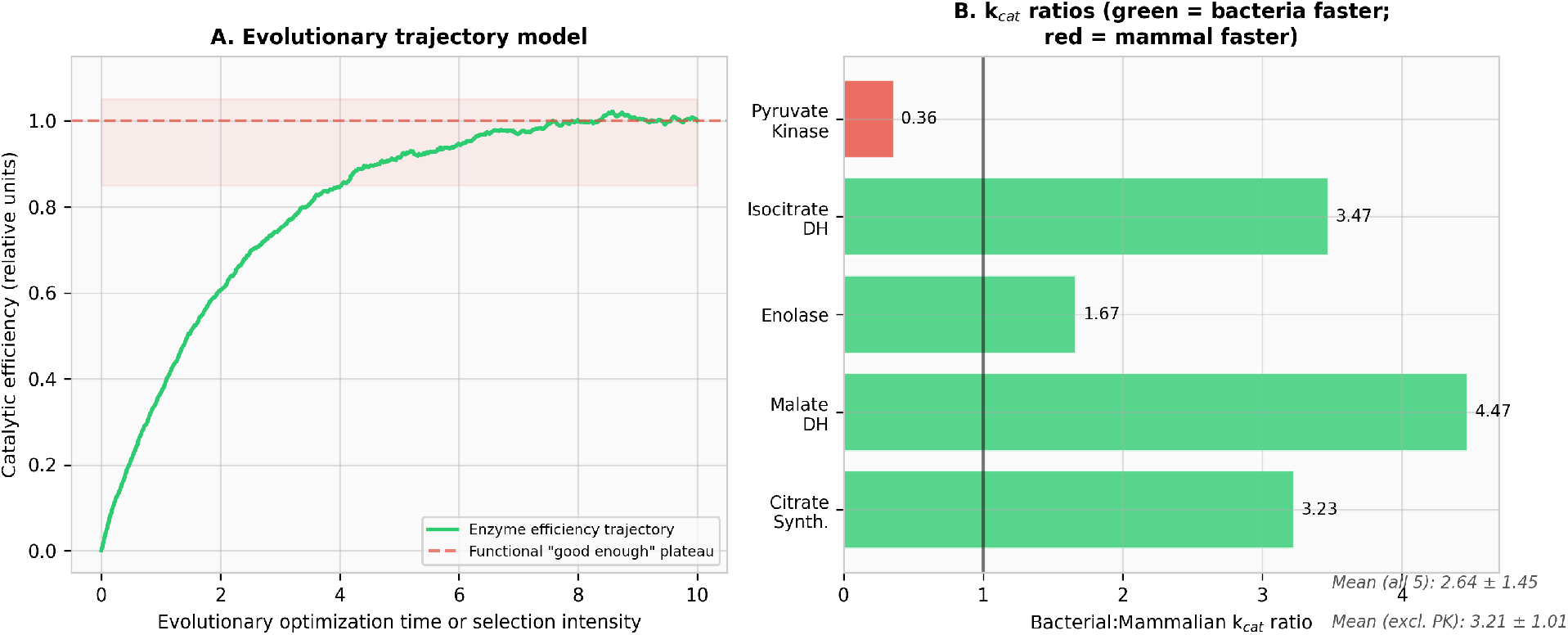
(A) Conceptual model of enzyme efficiency optimization: rapid initial rise followed by a “good enough” plateau, consistent with Labourel & Rajon (2021). (B) Bacterial:mammalian k_cat_ ratios for individual enzymes. Green = bacteria faster; red = mammal faster. The mean including all enzymes (2.64) differs substantially from the TCA-only mean (3.72).

### 3.3 Pathway-Specific Patterns

Grouping enzymes by metabolic pathway reveals distinct patterns (Figure 3). TCA cycle dehydrogenases consistently show higher bacterial k_cat_ (mean ratio 3.72×), reflecting the high constitutive TCA flux in aerobic bacteria. Glycolytic kinases and transferases show the opposite pattern (mean ratio 0.42×), driven by tissue-specific mammalian isoforms optimized for high glycolytic flux (e.g., RBC pyruvate kinase). OXPHOS complexes are remarkably conserved (mean ratio 1.1×), despite 2–4× more subunits in eukaryotic versions, indicating that added structural complexity serves regulatory rather than catalytic functions.

**Figure 3.**
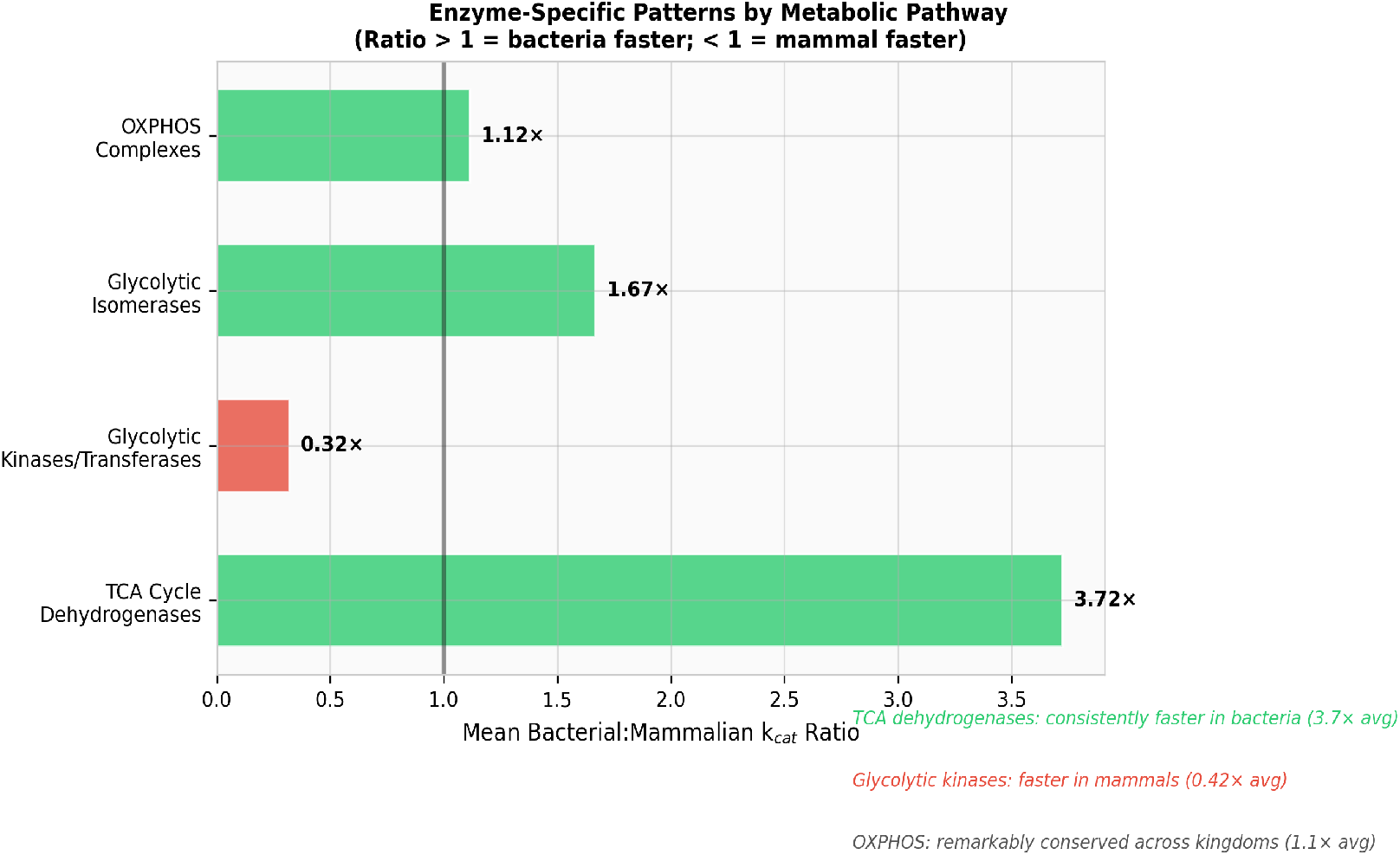
Enzyme-specific patterns by metabolic pathway. Mean bacterial:mammalian k_cat_ ratios grouped by pathway class. TCA dehydrogenases are consistently faster in bacteria; glycolytic kinases/transferases are faster in mammals; OXPHOS complexes are conserved.

### 3.4 Notable Exceptions and Counter-Examples

Four major counter-examples merit detailed discussion. (1) **Alkaline phosphatase**: Bovine intestinal AP isoforms (bIAP I: 1,800 s^−1^; bIAP IV: 6,100 s^−1^) are 30–100× faster than *E. coli* AP (∼60 s^−1^)^8,9^. (2) **Carbonic anhydrase**: Mammalian CA II achieves k_cat_ ∼10^6^ s^−1^, near diffusion-controlled^10^. (3) **Triosephosphate isomerase**: Both yeast and human TPI operate at k_cat_ ∼10^4^ s^−1^ near the diffusion limit, representing an evolutionary ceiling. (4) **GAPDH**: Yeast is ∼10× faster than *E. coli*, a eukaryote outpacing a prokaryote, driven by the Crabtree effect.

## 4. Discussion

### 4.1 Context-Specific Selection, Not Universal Complexity Trend

Our expanded analysis reveals that the relationship between organismal complexity and enzyme catalytic speed is **not a universal trend but a pathway-specific pattern** shaped by context-dependent selection pressures. The Bar-Even et al. (2011) analysis of ∼2,000 enzymes shows extensively overlapping k_cat_ distributions between prokaryotes and eukaryotes^2^, consistent with our finding that only specific enzyme classes (TCA dehydrogenases) exhibit the bacteria > mammal pattern.

We propose that the TCA dehydrogenase pattern reflects constitutively high TCA flux in aerobic bacteria, where these enzymes face strong selection for speed. In contrast, mammalian TCA enzymes operate under allosteric regulation (by ATP, NADH, Ca^2+^) that constrains maximal turnover in exchange for metabolic control^18^. The reversed pattern in glycolytic kinases reflects tissue-specific adaptation: RBCs and fermentative yeast require maximal glycolytic throughput, producing enzymes that exceed bacterial counterparts.

### 4.2 The “Rapid Bursts and Slow Declines” Framework

Newton et al. (2015) proposed that enzymes achieve near-peak catalytic performance in rapid evolutionary bursts when they become rate-limiting, followed by slow declines through neutral drift^3^. Ancestral enzyme reconstruction supports this: resurrected Precambrian thioredoxins were 150–200% more active than modern *E. coli* or human versions^19^. This suggests an alternative interpretation of our data: rather than bacteria “evolving faster enzymes,” eukaryotic TCA enzymes may have **decayed from ancestral peaks** through relaxed selection once cellular compartmentalization and metabolic channeling reduced the premium on individual enzyme speed.

### 4.3 OXPHOS Conservation and the Regulation vs. Speed Trade-off

The most striking finding is the near-identical catalytic rates of OXPHOS complexes across kingdoms despite dramatic structural elaboration. Mammalian ATP synthase (∼17 subunits) and *E. coli* ATP synthase (∼8 subunits) both operate at ∼270 s^−116^. This demonstrates that added eukaryotic subunits (the “stator stalk,” regulatory subunits, tissue-specific isoforms) serve regulatory and structural roles without altering core catalytic mechanism. The OXPHOS data powerfully illustrate that complexity and catalytic speed are orthogonal dimensions in enzyme evolution.

### 4.4 Practical Implications

These enzyme-specific patterns have direct implications for metabolic engineering and industrial biotechnology. When maximal flux is needed, bacterial TCA enzymes or yeast glycolytic enzymes provide faster catalytic chassis. When regulatory control is paramount, mammalian isoforms offer superior tunability. The OXPHOS conservation suggests that core catalytic mechanisms are optimized early and resistant to further improvement, regardless of organism.

## 5. Conclusions

Our investigation reveals that the catalytic efficiency of core metabolic enzymes does not diminish uniformly with organismal complexity. Instead, **pathway-specific patterns** emerge: TCA cycle dehydrogenases are 3–4× faster in bacteria, glycolytic kinases are often faster in mammals (in high-demand tissues), and OXPHOS complexes are remarkably conserved across kingdoms. These patterns reflect the intersection of metabolic demand, regulatory complexity, and evolutionary drift rather than a simple complexity gradient. The “rapid bursts and slow declines” framework provides the most parsimonious explanation: ancestral enzymes achieved near-peak performance, and subsequent evolution either maintained, relaxed, or redirected selection depending on metabolic context. Understanding these enzyme-specific patterns illuminates principles of molecular evolution and guides practical choices in metabolic engineering and enzyme sourcing.

## Acknowledgments

The authors acknowledge the BRENDA and BioNumbers databases for providing curated kinetic data.

## Declaration of Generative AI Use

During the preparation of this work, the authors used ChatGPT and Claude (Anthropic) for literature validation, data verification against primary sources, figure generation, and manuscript formatting. All content was reviewed and edited by the authors, who take full responsibility for accuracy and integrity.

